# Genetic parameters for resistance to gastrointestinal nematodes in sheep: a meta-analysis

**DOI:** 10.1101/2022.06.16.496403

**Authors:** Adam D. Hayward

## Abstract

Gastrointestinal nematodes (GIN) are damaging parasites of global sheep populations. The key weapons in fighting GIN have been anthelmintic drugs, but the emergence of drug-resistant parasites has meant that alternative control methods are needed. One of these alternatives is to breed for enhanced host resistance to GIN, and decades of research have estimated the genetic contribution to different measures of resistance to GIN and their genetic correlations with other desirable performance traits. It is clear that parasite resistance is a heritable trait that can be selected for. Despite this consensus, estimates of both heritability of resistance and genetic correlations with other traits vary widely between studies, and the reasons for this variation have not been examined. This study provides a comprehensive and quantitative meta-analysis of genetic parameters for resistance to GIN in sheep, including measures of worm burden (faecal egg counts, FEC), anti-parasite immunity (GIN-specific antibodies), and parasite-induced pathology (FAMACHA© scores). Analysis of 591 heritability estimates from 121 studies revealed a global heritability estimate for resistance to GIN of 0.25 (95%CI = 0.22 – 0.27) that was stable across breeds, ages, geographical location and analytical methods. Meanwhile, analysis of 559 genetic correlations from 54 studies revealed that resistance to GIN overall has a positive genetic correlation of +0.10 (95%CI = 0.02 – 0.19) with performance traits, and that this was consistent across breeds, ages, sexes and analytical methods. Importantly, the direction of the genetic correlation varied with the resistance trait measured: while FEC and FAMACHA© scores were favourably correlated with performance traits, adaptive immune markers were unfavourably correlated, suggesting that selection for enhanced immune responses to GIN could reduce animal performance. Overall, the results suggest that breeding for resistance to GIN should continue to form part of integrated management programs to reduce the impact of parasites on health and performance, but that selection for enhanced immune responses should be avoided.

## INTRODUCTION

Gastrointestinal nematodes (GIN) are among the most prevalent parasites of sheep worldwide and cause a significant economic impact on the industry through effects on animal production and the costs of treatments (Charlier et al., 2020). The most important GIN species in sheep in temperate regions are *Teladorsagia circumcincta, Trichostrongylus colubriformis* and *Nematodirus battus*, while the blood-feeding *Haemonchus contortus* is more dominant in the tropics. All of these species have direct life cycles: larvae are consumed during grazing and develop into adults in the gastrointestinal tract, which then shed eggs that are detectable in faeces. While the spatiotemporal epidemiology of the species varies somewhat – with *N. battus* being more common in lambs, and *H. contortus* more prevalent in tropical regions – coinfections are common. Control has typically focused on administration of various classes of anthelmintics, but resistance to these drugs is spreading among worm populations, leading to the call for more integrated parasite management (IPM) programs in the future (Kenyon et al., 2017).

One method of mitigating the impact of GIN in sheep populations that has been explored for decades is to use selective breeding to enhance the resistance of animals to infection. For selection to result in a response, the desired trait must have non-zero heritability (Falconer and Mackay, 1996). The heritability of a trait, *h^2^*, may be defined as the proportion of phenotypic variance in a trait due to additive genetic effects; a trait with *h^2^* = 0 has no additive genetic variance and a trait with *h^2^* = 1 is entirely determined by additive genetic effects. Selective breeding to mitigate the impact of GIN infection has largely been based on two traits: resistance, which is the ability of an animal to reduce parasite burden, and resilience, which is the ability of an animal to maintain performance in the face of infection (Bishop, 2012a). Resistance has been measured in various ways, including worm faecal egg counts (FEC), which estimate the host’s ability to limit worm number, size, or fecundity; immune markers such as nematode-specific antibodies or eosinophil counts, which are known to be actively involved in the acquired immune response to GIN (McRae et al., 2015); and FAMACHA© score, which is a marker of anaemia specifically used to monitor infection with the blood-feeding GIN *H. contortus* and the need for anthelmintic treatment (van Wyk and Bath, 2002). Of these traits, FEC has been subjected to the most research, partly because of its ease of measurement and its applicability to all of these nematode species. As has been outlined in many review articles (Bishop, 2012b; Bisset and Morris, 1996; Stear et al., 1997a; Stear et al., 2009; Zvinorova et al., 2016), nematode FEC has a considerable genetic component: it varies considerably among breeds and is a heritable trait within breeds. This heritable basis has been used to effectively breed low (and indeed high) FEC lines of sheep (Baker et al., 1990). Although subjected to less research, it is apparent that nematode-specific antibody responses (Murphy et al., 2010; Strain et al., 2002), eosinophil counts (Davies et al., 2005; Woolaston and Piper, 1996) and FAMACHA© scores (Álvarez et al., 2018; Cloete et al., 2016; Ngere et al., 2017) are also significantly heritable and hence could be used as target traits for selective breeding.

While it is firmly concluded that resistance to GIN is a heritable trait, qualitative reviews reveal that heritability estimates range 0-0.65 (Zvinorova et al., 2016), with typical values of 0.2-0.4 (Bishop, 2012b; Stear et al., 1997a). Similarly, heritability estimates of 0-0.4 have been reported for FAMACHA© scores (Ngere et al., 2017; Snyman and Fisher, 2019) and nematode-specific antibody responses (Davies et al., 2005; Gauly et al., 2002). Since accurately estimating the heritability of a given trait is essential for determining the likely response to a given (artificial or natural) selection pressure (Falconer and Mackay, 1996), it is important that variation in heritability estimates is understood. For example, many early studies estimated heritability using techniques such as parent-offspring regression or sibling analysis, while the “animal model” is the overwhelming choice currently; these variance-partitioning models fit the pedigree structure or genomic relatedness matrix as random effects and as such make fuller use of the relatedness or genomic data available for an individual (Henderson, 1950; Lynch and Walsh, 1998). Animal models generally report lower heritability estimates (Kruuk, 2004), and more recent studies may be based on larger sample sizes, but has this driven a change in estimates of the heritability resistance to GIN of over time? Estimates of the heritability of resistance to GIN in sheep have also been derived from animals of different breeds, sexes and ages, and in response to different GIN species under different environmental conditions. How do heritability estimates vary among different categories of animals infected with different parasite species? Lastly, which traits are likely to yield the greatest responses to selection? To address this question, it is essential to determine whether the heritability of resistance to GIN is dependent on the trait that is measured.

Resistance to GIN is a desirable trait to breed for because higher resistance (e.g. lower FEC) is often associated with improved animal performance (Cruz-Rojo et al., 2012; Fthenakis et al., 2015; Galyon et al., 2020; Idika et al., 2012; Mavrot et al., 2015; Notter et al., 2017). As such, it may be expected that breeding for increased resistance should also lead to improved performance. This may not, however, be the case, due to costs of resistance to infection, which include the investment of resources into immune responses rather than growth or reproduction; the negative effects of immunopathology; and potential trade-offs between immune responses to nematodes and other infections (Colditz, 2008). As such, we may expect to see negative genetic correlations between parasite resistance and performance traits such as body weight (Greer, 2008; Rauw et al., 1998). Genetic correlations are scaled from −1 (genes associated with high values of one trait are associated with low values of a second trait), through zero (genetic basis of two traits is unrelated), to 1 (genetic basis of two traits is identical). There is enormous variation in the strength of genetic correlations between resistance to GIN in sheep and performance traits, from highly unfavourable to highly favourable (Albers et al., 1987; Assenza et al., 2014; Bishop et al., 1996; Bishop et al., 2004; Brown and Fogarty, 2017; Douch et al., 1995a; Eady et al., 1998; Matebesi-Ranthimo et al., 2014; Shaw et al., 1999; Shaw et al., 2013; Stear et al., 2004; Wolf et al., 2008). Understanding the way in which aspects of study design influence the direction and strength of genetic correlations is therefore necessary to determine the correlated responses to selection for resistance on performance traits. In addition to those aspects described above, genetic correlations with resistance could vary depending on the performance trait being measured; such desirable traits could include body weight, body condition, or aspects of milk and wool quantity or quality.

Decades of empirical work estimating the heritability of resistance to GIN and its genetic correlations with other traits has concluded that resistance traits are heritable and that genetic correlations with performance traits vary. This work has been reviewed extensively, but qualitative reviews, however comprehensive, cannot address two key questions: first, what are the “average” estimates for the heritability of resistance to GIN and its genetic correlation with performance; second, what factors are likely to influence such estimates? In order to answer these questions, I employed meta-analysis, a set of statistical tools that enables the results of past research to be synthesized in a quantitative manner (Gurevitch et al., 2018; Koricheva et al., 2013; Lean et al., 2009; Nakagawa and Santos, 2012). Meta-analysis enables the estimation of “global” effects, which can be thought of as a weighted mean effect across all studies, and identify the impact of “moderator” variables such as study design on the results of past studies. While very common in medical science (Gurevitch et al., 2018; Lean et al., 2009), meta-analysis has rarely been seen in livestock science, although that picture has changed recently (Hayward et al., 2021; Mavrot et al., 2015; Zaatout and Hezil, 2022). Two recent meta-analyses have analysed published genetic parameters for a variety of economically-important traits in sheep, including body weight, growth, wool production, carcass parameters, reproductive parameters, longevity and diseases (Medrado et al., 2021; Mucha et al., 2022). These studies provided “global” heritability estimates for FEC and FAMACHA scores of 0.17 and 0.25 (Medrado et al., 2021) and estimates for FEC, parasite antibodies and parasite immunoglobulins of 0.29, 0.18 and 0.36 (Mucha et al., 2022). Furthermore, these studies reported a favourable “global” genetic correlation of −0.19 for FEC and body weight (Medrado et al., 2021) and non-significant correlations between FEC, parasite immunity and body weight (Mucha et al., 2022). Both studies reported substantial heterogeneity in genetic parameters, but neither investigated the moderator variables associated with this variation, and the breadth of economic parameters that were investigated did not facilitate an exhaustive search of the literature. Here, I used a comprehensive literature search and meta-analysis to first, estimate a global heritability specifically for resistance to GIN and its genetic correlation with performance, and second, to identify the impact of moderator variables on variation in study outcomes.

## METHODS

### Literature search and screening

I searched the scientific literature in order to find studies that had estimated the heritability of parasite resistance traits and/or genetic correlations between such traits and measures of animal performance. Searches of Web of Science, PubMed and Google Scholar were performed on April 1^st^ 2021 using the search terms (sheep OR lamb) AND (nematode OR strongyle OR Teladorsagia OR Trichostrongylus OR Haemonchus) AND (resistance OR faecal egg count OR antibody OR IgG OR IgA) AND (heritab* OR genetic). This search yielded 1810 items. To this, I added all literature cited by six influential review articles (Bishop, 2012b; Bisset and Morris, 1996; Stear et al., 1997a; Stear et al., 2001; Stear et al., 2009; Zvinorova et al., 2016) and all the literature cited in these articles using the Publish or Perish software (Harzing, 2007). These searches yielded another 1232 items. Finally, I searched the abstracts database of the Association for the Advancement of Animal Breeding and Genetics (http://www.aaabg.org/aaabghome/AllProceedings.php) with the terms “sheep, nematode, resistance”, yielding a further 402 items. The initial search therefore included a total of 3444 research items. I followed the PRISMA protocol for systematic review and meta-analysis (Moher et al., 2009) and provide a PRISMA checklist (**Table S1**) and flow diagram (**Figure S1**) to describe the search and filtering process in more detail.

I screened the titles and abstracts of these items and rejected 885 duplicate items (25.7%). I then removed a further 2181 items for a variety of reasons (**Table S2**), the most common of which were that they were studies on the wrong host species (13.7%), studies of anthelminthic resistance (13.4%), comparing resistant and susceptible genotypes of animal (9.0%), or investigated the wrong traits (7.2%). Of the initial 3444 items, 378 (14.8%) were considered for full review.

I included data from studies of sheep that estimated the heritability of traits associated with resistance to gastrointestinal nematodes including mixed-species infection, *T. circumcincta, T. colubriformis, N. battus* and *H. contortus*, and/or genetic correlations with traits including weight, weight gain, wool production, milk production, fatness, muscle depth, skeletal size, and reproductive output. 122/378 (32.3%) research items provided data that were suitable for inclusion, while 256 were excluded for a variety of reasons, including that they were review articles that provided no new data (22.5%), compared the performance of sheep selection lines (7.4%) and were studies of quantitative trait loci that did not report heritability estimates or genetic correlations (6.6%; **Table S3**). I recorded heritability estimates and/or genetic correlations, associated standard errors, and the number of samples that were used in the analysis. Where data were presented in figures but not in text or tables, I used the R package ‘metaDigitise’ to extract data (Pick et al., 2019). Once data screening and extraction were complete, the result was a heritability data set with 591 estimates from 121 sources and a genetic correlation data set of 559 estimates from 54 sources (**Table S4**).

### Data synthesis

Heritability and genetic correlation estimates came from studies that varied widely in a number of parameters including those related to the animals (breed, location, age, sex), research methods (infection type, statistical analysis), measures of resistance and gastro-intestinal nematode species (**Table S4**). Since heritability estimates and genetic correlations are not normally distributed, we used Fisher’s z-transformation (0.5(ln((1 + h^2^)/(1 − h^2^)))) before all analyses and back-transformed these effect sizes (*Zr*) to heritabilities for interpretation. Our values of *Zr* for both heritability and genetic correlation data sets approximated a normal distribution, and this method best adhered to the requirements of meta-analysis and was previously used in other similar meta-analyses (Bell et al., 2009; Dochtermann et al., 2019). The sampling variance of *Zr* is 1/(n − 3), where *n* is the number of individuals in the study. As such, heritability and genetic correlation estimates based on a larger number of individuals have a lower sampling variance.

The heritability data came from 121 sources (1-24 estimates per source), with data collected from 32 breeds of sheep, plus crosses, which were combined to a single breed. Host age was represented by a categorical variable where age was “lamb” (≤52 weeks, 76%), “adult” (>52 weeks, 12%) or “mixed” (12%). The heritability estimates were largely from mixed-sex groups of animals (84%), but there were also a number of estimates derived purely from females (12%) or males (4%). 82% of our estimate came from studies involving natural infections, while 18% were from studies where animals were experimentally infected. Studies also varied in statistical methods: while 77% of the estimates were from “animal models” using either pedigrees or genomic relatedness, the remaining 23% used methods including parent-offspring regression, sib analysis and sire models; all of these different methods were grouped to give a variable with two levels, namely “animal model” and “other”. Some studies considered maternal effects in their models: those that explicitly accounted for maternal effects, or stated that they did so in preliminary analysis but found them to be zero, were grouped together (27%), while studies that did not account for maternal effects were grouped together (73%). Host were infected with a variety of different nematode species: 49% did not specify the predominant parasite species and so we simply denoted them as “strongyle”, while others explicitly mentioned that *Haemonchus contortus, Teladorsagia circumcincta, Trichostrongylus colubriformis* or *Nematodirus* spp were dominant in (natural) multi-species infections (23%, 7%, 14% and 7% respectively). 3 estimates from 2 studies specifically mentioning *Cooperia* spp and *Strongyloides* spp were incorporated into the “strongyle” group. Finally, our heritability estimates were from a variety of traits related to parasite resistance: the majority of these were of faecal egg counts (FEC, 73%), but there were a number of other measures. 19% represented heritability estimates for immune traits; these included parasite-specific antibody responses from a variety of isotypes and specificities, as well as eosinophil counts (8% of the immune measures), while 5% were from FAMACHA© scores associated with *H. contortus* infection and 3% were from worm-specific traits, including worm number, length and fecundity. A breakdown of these variables is shown in **Figure S2**.

The genetic correlation data were from 54 sources (1-72 estimates per source), with data collected from 20 breeds of sheep, including crosses. 73% of these records were from lambs, with 20% from adults and 7% from mixed-aged groups; age in weeks was available for 362 data points. Genetic correlations were largely from mixed-sex groups (78%), with 18% from females and 4% from males. 86% of the genetic correlations were from natural infections and 14% were from experimental infections. 83% of studies came from multivariate animal models and 17% used other methods, while 38% accounted for maternal effects and 62% did not. Among parasite taxa, 54% of genetic correlations were assigned as ‘Strongyle’ (usually natural infections), while 21% were *Haemonchus contortus*, 1% were *Teladorsagia circumcincta*, 16% were *Trichostrongylus colubriformis* and 8% were *Nematodirus* spp. Of the parasite resistance traits, there were no genetic correlations involving worm traits, but 78% of correlations involved FEC, 18% immune measures and 4% FAMACHA© scores. Importantly, genetic correlations involving FEC and FAMACHA© scores were reversed before analysis, such that positive correlations were favourable.

Finally, genetic correlations were estimated between resistance traits and a variety of performance measures as follows: (1) any trait that was a weight measured at a single time point was considered to be a “weight” trait (38%); (2) any traits relate to fleece characteristics, including fleece weights, staple length or growth were grouped as “wool” traits (26%); (3) body condition score, fat depth, and muscle depth were grouped together as “condition” (18%); (4) weight gain of an individual measured between two time points, generally given as daily weight gain, was assigned as “DWG” (10%); (5) maternal traits such as lambs born, lamb weight and lamb growth rates were assigned as “reproductive” traits (5%); (6) 2% were traits related to milk production and quality; and (7) 1% were traits related to skeletal size such as leg length. A breakdown of these variables is shown in **Figure S3**.

### Meta-analysis

All analyses were performed in R version 4.1.2 (R Core Team, 2021). Meta-analysis was performed using the *rma.mv(*) function from the R package ‘metafor’ (Viechtbauer, 2010) version 3.0-2 and orchard plots were created using the package ‘orchaRd’ (Noble, 2021) version 0.0.0.9000. Separate meta-analyses were applied to the heritability and genetic correlation data sets.

First, global estimates of the heritability of nematode resistance and its genetic correlation with performance were gained using separate multi-level analyses. In order to account for non-independence of effect sizes derived from the same studies and from sheep of the same breed, random-effects models were fitted with the random effects of study and breed, plus an observation-level random effect (Viechtbauer, 2010) in order to estimate the residual variance. We tested the significance of random effects using likelihood ratio tests (LRTs), but both random effects were retained even if not supported in order to account for all of the variation when estimating the global effect. The heterogeneity in effect sizes was quantified by calculating the proportion of total variance that was due to variation in effect sizes (the *I^2^* statistic), where the remainder is accounted for by sampling error. Since random effects were fitted to account for similarity between effects from the same studies and breeds, modified *I^2^* was calculated for the between-study, between-breed, and residual effects (Nakagawa and Santos, 2012).

Next, a meta-regression approach was used in order to identify moderator variables that influenced the size of heritability and genetic correlation estimates. The absolute latitude that the study took place at was accounted for: this was either given in the paper, or if not, estimated based on the description of the study location, or the host institute of the lead author of the paper. Other moderators included the sex of the animals (female, mixed, or male); age category (lamb, mixed, or adult); infection type (natural or challenge); analytical method (animal model or other); whether maternal effects were accounted for (yes or no); the resistance trait being analysed (worm-specific trait, immune trait, FEC or FAMACHA© score); the parasite species (‘Strongyle’, *‘Haemonchus’, ‘Nematodirus’, ‘Teladorsagia’, ‘Trichostrongylus’*). For analysis of genetic correlations, we also included a moderator variable specifying the performance trait that was measured, which was a categorical variable that distinguished studies of weight, daily weight gain, body condition (including fat and muscle depth), wool production, milk production, skeletal size, and reproductive performance.

We also included two further moderators in order to account for publication bias in our model (Nakagawa et al., 2022). First, we included the square root of the sampling variance (*se_i_*) in order to account for bias caused by the tendency to publish only significant results; a significant positive effect would indicate that studies with higher sampling variance (i.e. high uncertainty) are only published when the heritability estimates are high. Second, we included the year of publication in order to test for a predicted decline in heritability estimates over time generated by factors that could include more of a bias towards publishing significant results earlier in a topic’s history (Koricheva and Kulinskaya, 2019), improved estimation methods, and larger sample sizes.

All moderator variables were fitted in the same meta-regression model and we determined whether the moderator was supported (i.e. whether the effect size varied according to the moderator) using Wald-type χ^2^ tests (*QM*). Where a categorical variable with more than two levels was deemed significant based on *Q_M_*, we conducted post-hoc tests to determine which levels of the variable differed from each other using the glht() function in the package ‘multcomp’ (Hothorn et al., 2008) with Benjamini-Hochberg correction. We also examined whether each level within categorical moderators was significantly different from the reference category using z-tests. In all analyses, P values ≤0.05 were deemed to be statistically significant.

## RESULTS

### Heritability of resistance to gastrointestinal nematodes

Multi-level meta-analysis of 591 heritability estimates from 121 studies provided a global estimate for the heritability of resistance to gastrointestinal nematode infection *β_global_*) of 0.2461 (95%CI = 0.22 – 0.27, z = 18.50, P<0.001; **Figure 1A**). Total heterogeneity was high (*I^2^* = 98.25%), with substantial heterogeneity accounted for by the study (*I^2^* = 30.09%, likelihood ratio test *χ^2^* = 45.48, P<0.001), but little variation between breeds (*I^2^* = 3.17%, *χ^2^* = 0.21, P=0.644) and with the remainder accounted for by residual effects (*I^2^* = 65.00).

**Figure 1.**
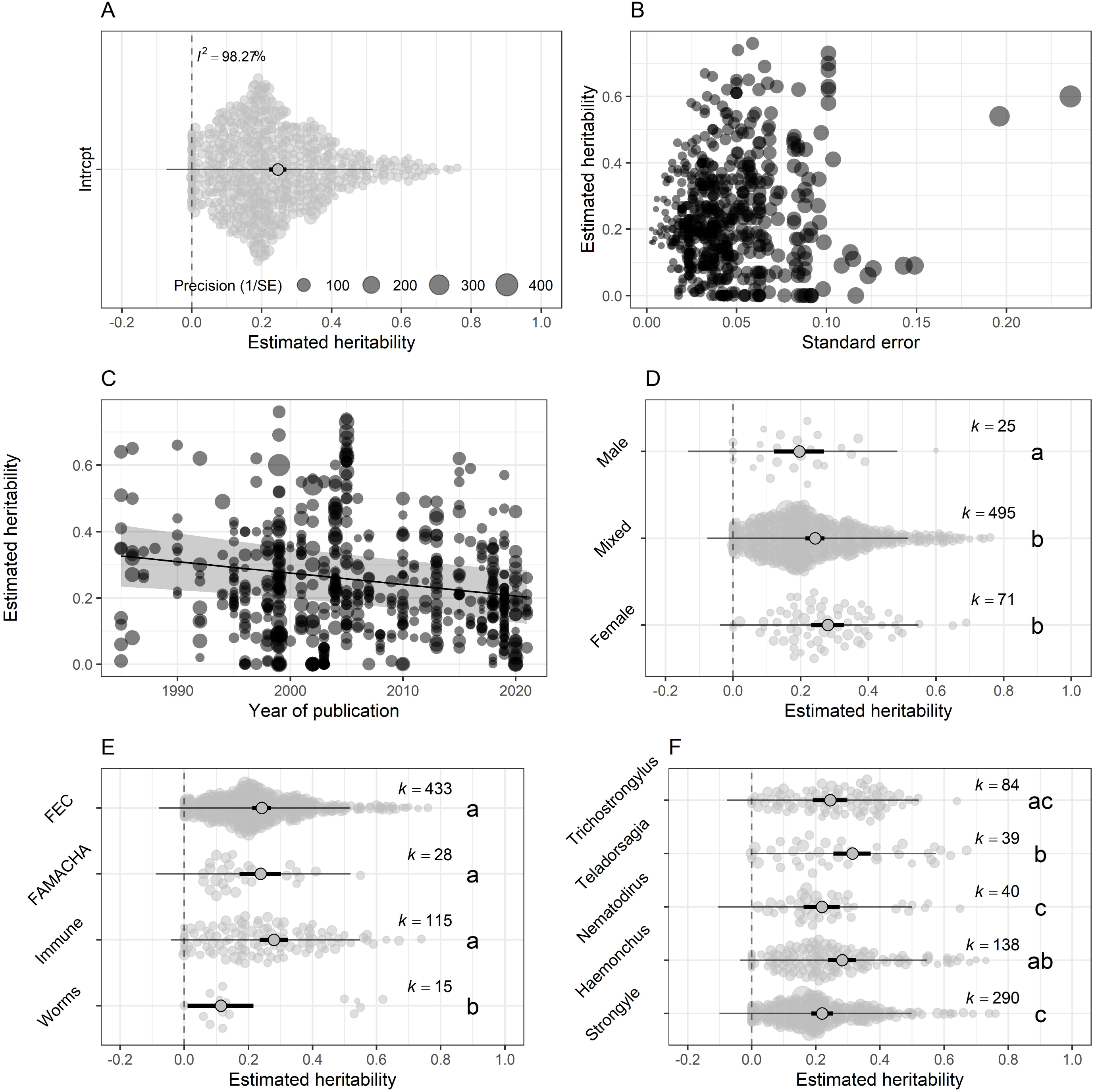
Orchard plots showing the results of meta-analysis of heritability of resistance to gastrointestinal nematodes in sheep. (A) shows the result of the random-effects model estimating the global heritability estimate, while the remainder show the results of the meta-regression model estimating the effects of moderators: (B) *se_i_* (not statistically significant); (C) year; (D) sex; (E) resistance trait; (F) parasite species. In (B) and (C), points show individual heritability estimates, with larger points representing studies with larger *se_i_* in (C) the regression line and shaded area indicate the predictions from the meta-regression. In (A) and (D-F), central circles show the estimate from the meta-regression; thick black lines show the 95%CI associated with the estimate; thin black lines show 95% prediction intervals (where 95% of the estimates lie) and each point shows an individual heritability estimate, sized based on the precision. Points are jittered in A and D-F in order to better display the spread of data. Lower-case letters (a, b, c) indicate differences between levels of categorical variables based on post-hoc tests (see Methods for details); *k* shows the number of data points available for each level of the categorical moderators.

Meta-regression analysis revealed that heritability estimates were not significantly influenced by latitude, age, infection type, statistical method, or maternal effects (**Table 1**). Heritability estimates were, however, influenced by the sex of the animals studied (*Q_M_*=6.66, DF=2, P=0.036); specifically, heritability estimates in studies of males were lower than those derived from studies of animals of mixed sex and females (**Figure 1D**). Heritability estimates were also influenced by the parasite resistance trait that was analysed (*Q_M_*=12.40, DF=3, P=0.006), with heritability estimates for worm-specific traits overall only just above zero, while those for FEC, FAMACHA© and immune traits were all >0.2 (**Figure 1E**). Heritability estimates also varied between parasite species (*Q_M_*=18.22, DF=4, P=0.001); post-hoc comparisons revealed that heritability estimates from infections with strongyles, *Nematodirus* and *Trichostrongylus* tended to be lower (~0.2) than those for *Haemonchus* and *Teladorsagia* predominant infections (~0.3; **Figure 1F**). Despite these differences, the overall heritability estimates for all parasites were comfortably above zero.

**Table 1.** Results from meta-regression of 591 estimates from 121 studies of the heritability of resistance to gastrointestinal nematodes. *Q_M_* and degrees of freedom (DF) and P values refer to the test for significant differences in effect size due to a moderator, while z and P values refer to tests for differences from the reference category. In each case, moderator estimates are relative to the reference category.

| Variable | Level | $Q_M$ | DF | P | Estimate | L-95%CI | U-95% CI | Z | P |
| --- | --- | --- | --- | --- | --- | --- | --- | --- | --- |
| Intercept | Intercept |  |  |  | 0.0163 | -0.1308 | 0.1634 | 0.22 | 0.828 |
| $se_i$ | $se_i$ | 3.6 | 1 | 0.058 | 0.7677 | -0.0257 | 1.5611 | 1.90 | 0.058 |
| Year | Year | 7.42 | 1 | 0.007 | -0.0035 | -0.0060 | -0.0010 | -2.72 | 0.007 |
| Latitude | Latitude | 3.56 | 1 | 0.059 | 0.0016 | -0.0001 | 0.0034 | 1.89 | 0.059 |
| Sex | Female | 6.66 | 2 | 0.036 |  |  |  |  |  |
|  | Mixed |  |  |  | -0.0118 | -0.0646 | 0.0411 | -0.44 | 0.662 |
|  | Male |  |  |  | -0.0995 | -0.1765 | -0.0225 | -2.53 | 0.011 |
| Age category | Lamb | 1.52 | 2 | 0.469 |  |  |  |  |  |
|  | Mixed |  |  |  | -0.0386 | -0.1002 | 0.0229 | -1.23 | 0.219 |
|  | Adult |  |  |  | -0.0036 | -0.0503 | 0.0430 | -0.15 | 0.878 |
| Infection type | Challenge | 0.86 | 1 | 0.354 |  |  |  |  |  |
|  | Natural |  |  |  | 0.0322 | -0.0359 | 0.1002 | 0.93 | 0.354 |
| Method | Animal model | 0.17 | 1 | 0.677 |  |  |  |  |  |
|  | Other |  |  |  | 0.0118 | -0.0436 | 0.0672 | 0.42 | 0.677 |
| Maternal | No | 3.26 | 1 | 0.071 |  |  |  |  |  |
|  | Yes |  |  |  | -0.0442 | -0.0921 | 0.0038 | -1.81 | 0.071 |
| Trait | Worms | 12.4 | 3 | 0.006 |  |  |  |  |  |
|  | Immune |  |  |  | 0.1612 | 0.0613 | 0.2610 | 3.16 | 0.002 |
|  | FAMACHA© |  |  |  | 0.1845 | 0.0633 | 0.3056 | 2.98 | 0.003 |
|  | FEC |  |  |  | 0.1857 | 0.0817 | 0.2898 | 3.50 | 0.001 |
| Parasite | Strongyle | 18.22 | 4 | 0.001 |  |  |  |  |  |
|  | *Haemonchus* |  |  |  | 0.0886 | 0.0279 | 0.1493 | 2.86 | 0.004 |
|  | *Nematodirus* |  |  |  | -0.0073 | -0.0606 | 0.0461 | -0.27 | 0.790 |
|  | *Teladorsagia* |  |  |  | 0.1236 | 0.0513 | 0.1960 | 3.35 | 0.001 |
|  | *Trichostrongylus* |  |  |  | 0.0308 | -0.0362 | 0.0979 | 0.90 | 0.368 |

We tested for publication bias through including the square root of the sampling variance (*se_i_*) in our meta-regression, and we found no support for an effect of *se_i_* (**Figure 1B**), although there was a tendency for increased *se_i_* to be associated with a higher heritability estimate (estimate = 0.7677, 95%CI = −0.0257 – 1.5611, *Qm*=3.60, DF=1, P=0.059), suggesting that published studies with lower sample sizes tended to have higher heritability estimates. We also tested for publication bias by including the effect of publication year, which we did find statistical support for (estimate = −0.0035, 95%CI = −0.0060 – −0.0010, *Q_m_*=7.42, DF=1, P=0.007), which suggested that heritability estimates have reduced in size with time (**Figure 1C**). By comparing estimates from models with and without the moderators of *se_i_* and year (Nakagawa et al., 2022), we found that the effect of this publication bias on the overall estimate of heritability was minimal: without *se_i_* and year our estimate was 0.25 (95%CI = 0.21-0.28), and with them it was 0.22 (95%CI = 0.19-0.26).

### Genetic correlations between parasite resistance and performance

Multi-level meta-analysis of 559 genetic correlations from 54 studies provided a global estimate (*β_global_*) of +0.1020 (95%CI = 0.0154 – 0.1871, z = 2.31, P=0.021; **Figure 2A**). Overall, this suggests a positive (i.e. favourable) genetic correlation between gastrointestinal nematode resistance and performance in sheep. Total heterogeneity was very high (*I^2^* = 99.78%), with substantial heterogeneity accounted for by the study (*I^2^* = 49.60%, *I^2^* = 174.71, P<0.001), little variation between breeds (*P* = 4.86%, *χ^2^* = 1.38, P=0.241) and the remainder accounted for by residual effects (*I^2^* = 45.43).

**Figure 2.**
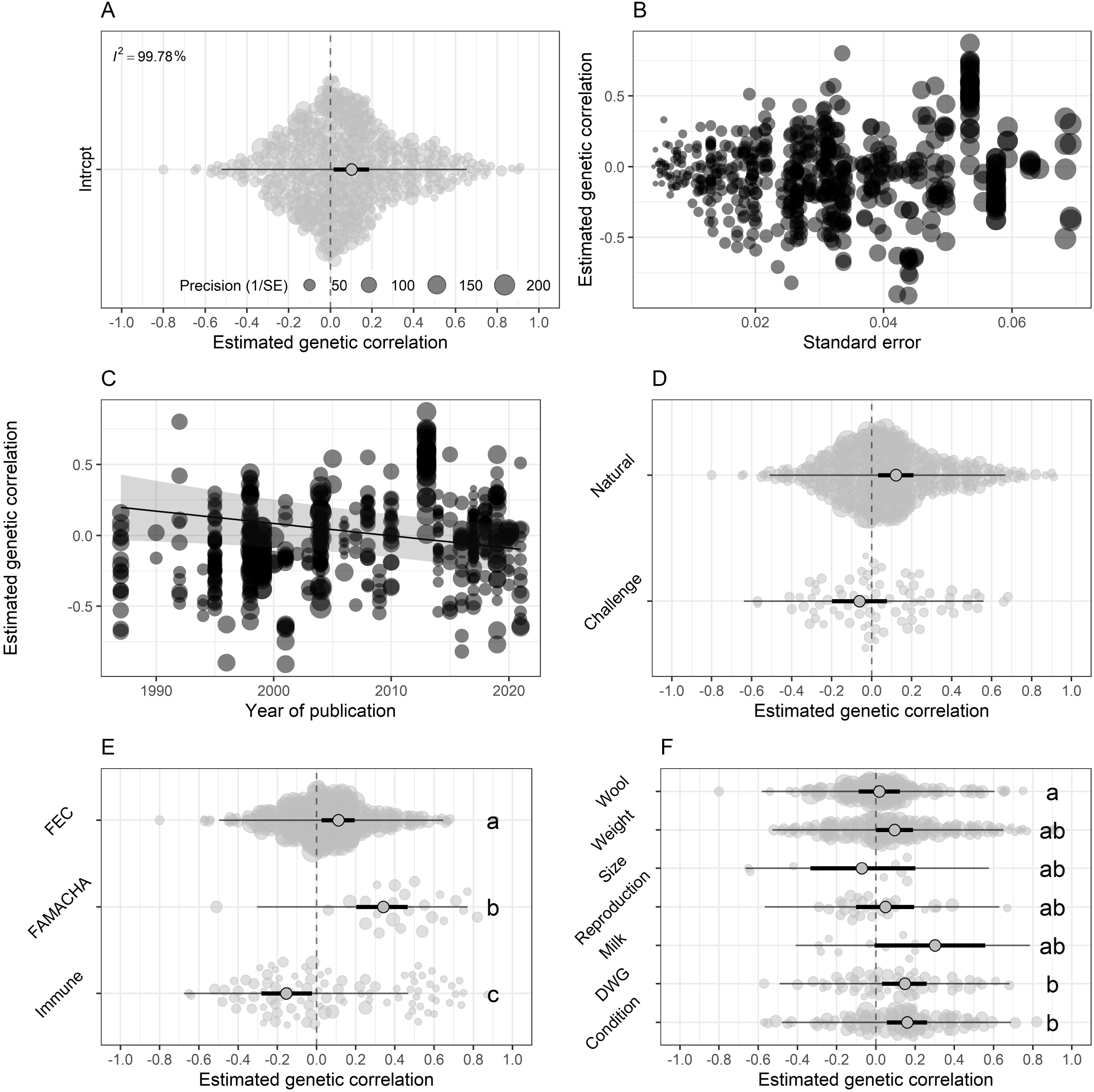
Orchard plots showing the results of meta-analysis of genetic correlations between resistance to gastrointestinal nematodes and performance in sheep. (A) shows the result of the random-effects model estimating the global genetic correlation estimate, while the remainder show the results of the meta-regression model estimating the effects of moderators: (B) *se_i_* (not statistically significant); (C) year; (D) infection method; (E) resistance trait; (F) performance trait. In (B) and (C), points show individual genetic correlation estimates, with larger points representing studies with larger *se_i_*; in (C) the regression line and shaded area indicate the predictions from the meta-regression. In (A) and (D-F), central circles show the estimate from the meta-regression; thick black lines show the 95%CI associated with the estimate; thin black lines show 95% prediction intervals (where 95% of the estimates lie) and each point shows an individual heritability estimate, sized based on the precision. Lower-case letters (a, b, c) indicate differences between levels of categorical variables based on post-hoc tests (see Methods for details).

Meta-regression analysis revealed that genetic correlations between parasite resistance and performance were not significantly influenced by latitude, age, sex, statistical method, maternal effects, or parasite species (**Table 2**). Genetic correlations were, however, influenced by the type of infection that was used in the study (*Q_M_*=4.15, DF=1, P=0.042), with genetic correlations more strongly positive under natural infection and not differing from zero under experimental challenge (**Figure 2D**). The trait used to assess parasite resistance was also important in determining the strength and direction of the genetic correlation with performance (*Q_M_*=31.46, DF=2, P<0.001): results suggested that genetic correlations involving FEC were around the global estimate of approximately +0.1, those involving FAMACHA© scores were more strongly positive (around +0.3), but those involving immune markers were negative (around −0.15; **Figure 2E**). Finally, the strength of genetic correlations also depended on the performance measure that was used (*Q_M_*=19.94, DF=6, P=0.003). Specifically, *post-hoc* analysis revealed that genetic correlations between parasite resistance and wool traits were lower than those between parasite resistance and both DWG and condition, with all other traits being statistically indistinguishable from every other trait (**Figure 2F**).

**Table 2.**
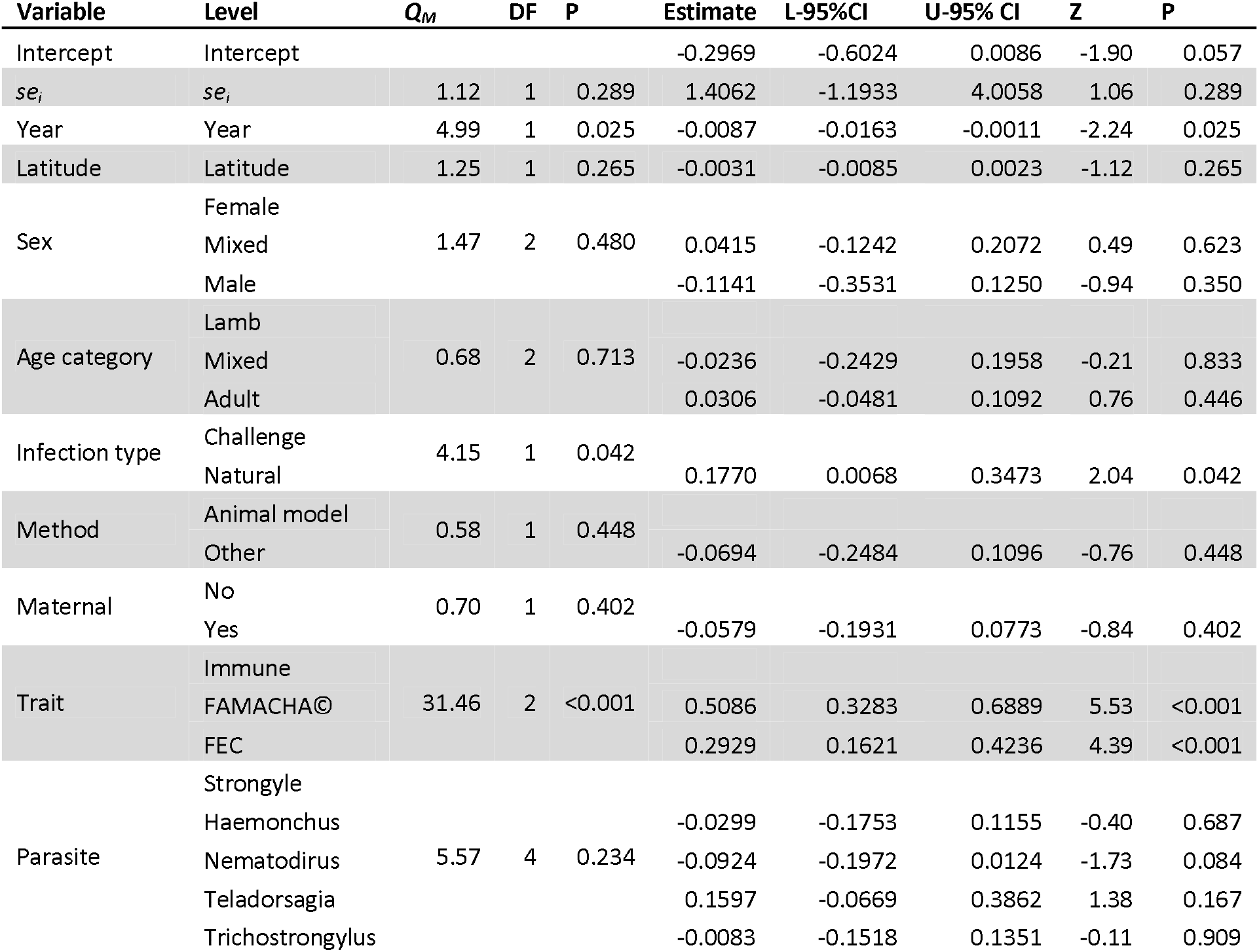
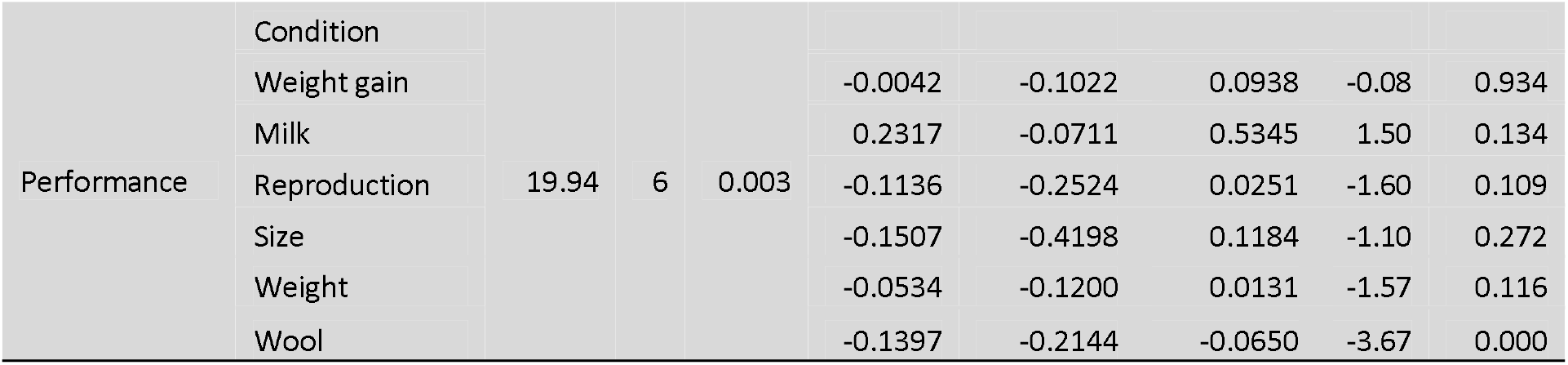
Results from meta-regression of 559 estimates from 54 studies of genetic correlations between resistance to gastrointestinal nematodes and performance in sheep. *Q_M_* and degrees of freedom (DF) and P values refer to the test for significant differences in effect size due to a moderator, while z and P values refer to tests for differences from the reference category. In each case, moderator estimates are relative to the reference category.

Analysis of publication bias through meta-regression did not support an effect of *se_i_*(estimate = 1.4062, 95%CI = −1.1933 – 4.0058, *Qm*=1.12, DF=1, P=0.289), suggesting that the strength and direction of published genetic correlations did not depend on sample size (**Figure 2B**). There was, however, support for the effect of year (estimate = −0.0087, 95%CI = −0.0163 – −0.0011, *Q_m_*=4.99, DF=1, P=0.025), which suggested that there has been a trend for genetic correlations to move from strongly positive to neutral between 1980s and the present (**Figure 2C**). As with heritability estimates, the impact of these sources of publication bias on the results were minimal: estimates from the model including *se_i_* and year were only marginally stronger (+0.0231) compared to a model without them.

## DISCUSSION

Gastrointestinal nematodes (GIN) are among the most prevalent and damaging parasites infecting domestic sheep populations worldwide. The possibility of breeding to enhance resistance to GIN has been subjected to decades of research, and findings suggest that resistance is underpinned by appreciable genetic variation (Bishop, 2012b; Bisset and Morris, 1996; Stear et al., 1997a). Estimates of the heritability of resistance to GIN vary widely, however, and it has yet to be determined which aspects of study design influence this variation in results. Furthermore, a potential drawback to breeding for resistance is the potential for negative genetic correlations with other desirable traits (Greer, 2008), but once again, genetic correlations between resistance and performance vary widely and the factors underpinning this variation are not understood. Here, a meta-analytic approach has addressed these knowledge gaps, providing global estimates for the heritability of resistance to GIN in sheep and its genetic correlation with performance. Our results also show that variation in heritability estimates is influenced by the parasite and resistance traits measured, and even more strikingly, genetic correlations between resistance and performance vary for different aspects of parasite resistance. Finally, both heritability and genetic correlations for parasite resistance have changed across time.

Published estimates of the heritability of resistance to GIN vary widely and have been the subject of a large number of review articles (Amarante and Amarante, 2003; Barger, 1993; Bishop, 2012b; Bishop and Morris, 2007; Karrow et al., 2014; Saddiqi et al., 2011; Sayers and Sweeney, 2005; Stear et al., 2001; Stear et al., 2009; Stear and Murray, 1994; Venturina et al., 2013; Windon, 1996) and they often describe resistance to GIN as ‘moderately heritable’ and provide a range of estimates of around 0.2-0.4. Our global estimate for the heritability of parasite resistance (0.25, 95%CI = 0.22 – 0.27) fell within this range, confirming the appreciable heritability of resistance to GIN in sheep. This estimate is also broadly in agreement with previous studies that have attempted to synthesise research on genetic parameters for resistance to GIN in sheep. These include a weighted mean estimate of the heritability of GIN FEC of 0.27±0.02 (Safari et al., 2005), although this was calculated from just 16 heritability estimates. Our estimate is also broadly comparable to those of a previous meta-analysis of genetic parameters for economic traits in sheep, which reported global heritability estimates of 0.17 (95%CI = 0.15 – 0.19) for FEC and 0.26 (95%CI = 0.19 – 0.32) for FAMACHA© scores (Medrado et al., 2021), although our estimate for FEC is slightly higher. This recent study, however, only assessed estimates from 37 articles for FEC and 11 for FAMACHA©, and their literature search only returned 505 heritability estimates for 35 traits including FEC and FAMACHA, compared to the search in the present study, which returned 591 estimates for parasite resistance traits alone. Another study provided a meta-analysis of “resilience traits” in sheep and goats using data derived from 13 EU partners and published articles (Mucha et al., 2022). These included heritability estimates of 0.29±0.03SE for FEC, 0.18±0.07 for “parasite antibody” and 0.36±0.06 for “parasite immunoglobulin”, based on 116, 6 and 24 estimates respectively, although the literature search process was not transparent and the difference between parasite “antibody” and “immunoglobulin” traits was unclear. Despite these previous studies, the present study is the most comprehensive synthesis of results on genetic parameters for resistance to GIN yet undertaken, reviewing the largest number of studies and unlike previous studies, has studied the influence of moderators on heritability estimates.

Some studies have found variation in heritability estimates among sheep of different ages (Baker et al., 2003; Brown et al., 2013; Roy et al., 2018; Snyman and Fisher, 2019; Stear et al., 2004) and with different estimation methods (Ngere et al., 2018), and it has also been observed that ‘animal models’ can return lower heritability estimates than other methods because they can, for example, account for common environment effects (Kruuk, 2004). Interestingly, meta-regression analysis revealed that neither sheep age, nor the statistical method used, had a consistent impact on the global heritability estimate. In contrast, we found that sex was associated with variation in heritability estimates, with heritability lower in studies of purely males (**Table 1**). This difference was based on a small sample size, however, with only 25 estimates coming from males (**Figure 1D**) out of the 591 estimates analysed. Nevertheless, these differences could have resulted from sex-specific genetic architecture of parasite resistance in sheep, or potentially a larger contribution of environmental or “residual” effects in males compared to the other groups. We also found variation among the resistance trait analysed, with worm-specific traits having an overall lower heritability estimate compared to FEC, FAMACHA© and immune traits (**Table 1**). In this case, the difference was substantial, with the 95%CI of the heritability estimate for worm-specific traits barely above zero (**Figure 1E**), but these estimates were based on just 15 heritability estimates from 3 studies. By examining these studies more closely, it is apparent that traits related to the number of worms tended to have low heritability estimates, while those associated with the size and fecundity of the worms, which are themselves linked, albeit in a curvilinear fashion (Stear and Bishop, 1999), tended to have higher heritability estimates. For example, one study of Scottish Blackface lambs found that the heritability of the numbers of nematode larval stages (L4 and L5) and adult worms varied between 0 and 0.14, while the number of eggs *in utero* in female worms and worm length had heritabilities of 0.55 and 0.62 respectively (Stear et al., 1997b). In a similar study of Scottish Blackface lambs, heritability for counts of worms of various stages varied between 0.06 and 0.13, while eggs *in utero* and length again had much higher values of 0.50 and 0.53 (Davies et al., 2005). It therefore seems that host genetic control is stronger over worm length and fecundity, rather than worm number, which is consistent with antibody-mediated reduction in worm size rather than numbers during the acquired immune response to GIN (Halliday et al., 2007; McRae et al., 2015; Venturina et al., 2013). Finally, heritability estimates varied depending on the worm species (**Table 1**), although differences between parasite species were small and all heritability estimates were 0.2-0.3 (**Figure 1F**).

The nature of the genetic relationship between resistance to GIN and desirable production traits such as weight, growth, and milk and wool production is crucial, since it determines the likely consequences of selection for resistance for other traits. Our meta-analysis of 559 genetic correlations from 54 studies suggested that, overall, there was a relatively weak but statistically significant positive genetic correlation of +0.10 (95%CI = 0.02 – 0.19) between resistance to GIN and sheep performance (**Figure 2A**). A previous meta-analysis suggested a stronger genetic correlation between GIN FEC and body weight, estimating this to be −0.19 (95%CI = −0.27 − −0.11; (Medrado et al., 2021)); note that in this case, the correlation was not reversed, so this is equivalent to an estimate of +0.19 in the present study. Once again, this was based on data from only 19 studies, but adds to the narrative that genetic correlations between parasite resistance and performance are not unfavourable.

The global estimate presented in the present study suggests that, very broadly, selection of animals for increased parasite resistance should enhance other aspects of performance, and that it should certainly not be traded-off against performance as has been feared (Greer, 2008; Rauw et al., 1998). There is much subtlety underpinning this, however. For example, I found that genetic correlations between resistance and performance were generally more strongly positive in natural infections than in experimental infections, potentially because experimental challenges are often sub-clinical. Furthermore, the strength and direction of genetic correlations between resistance to GIN and animal performance depended on the resistance and performance traits studied. In particular, while FEC and FAMACHA© scores were favourably correlated with performance traits, immune markers were negatively associated with performance at the genetic level (**Figure 2E**). That FAMACHA© scores are positively genetically correlated with parasite resistance is expected, given that FAMACHA© is a marker of the anaemia caused by *Haemonchus*, and as such encompasses not only the parasite burden (resistance) but also direct damage caused by the parasite (van Wyk and Bath, 2002), which may be influenced by the tolerance of the host and/or the virulence of the parasite. It is logical to expect that anaemic hosts should be lighter as this indicates blood feeding by the parasite and associated loss of circulating nutrients to the host, and indeed only one genetic correlation involving FAMACHA© score and performance included in this meta-analysis was negative, derived from a study where there was a positive genetic correlation between FAMACHA© score and body weight, such that animals with more anaemia were heavier (Rodrigues et al., 2021). More interestingly, and with great relevance to the idea that animal can be bred based on their immune responses to infection, was the finding that the genetic correlation between markers of immunity and animal performance was overall estimated to be negative (**Figure 2F**). A trade-off between immune responses and aspects of performance such as weight gain and milk production could exist for a number of reasons: costs of immunity include resource costs, immunopathology costs and costs of mounting responses to parasites that elicit contrasting types of immune response (Colditz, 2008). The results presented here therefore suggest that the hope that markers such as circulating levels of parasite-specific antibodies, although useful for selecting animals resistant to infection, may actually be counter-productive in terms of animal performance (Douch et al., 1995a; Douch et al., 1995b; Shaw et al., 1999). On the other hand, one immune marker was consistently positively associated with performance at the genetic level: the antibody response to *Trichostrongylus colubriformis* L3 larvae-specific carbohydrate larval antigen (CarLA) measured in saliva samples, which has been suggested as a suitable non-invasive measure of resistance (Shaw et al., 2012). This marker, which is heritable, consistently had genetic correlations with body weight, fleece weight, and weight gain, with 31/36 estimates greater than +0.30 (Shaw et al., 2013): it is also negatively correlated with worm FEC (Borkowski et al., 2020). Although these estimates all came from a single study, they are promising and deserve to be replicated more widely in order to validate salivary CarLA antibodies as a useful marker of resistance to infection that is favourably associated with animal performance.

I assessed publication bias by including the square root of the sampling variance (*se_i_*) in our meta-regression, in order to determine whether studies with larger error variance produce larger estimate, and year of publication, in order to test for changes in the size of reported results across time (Nakagawa et al., 2022). There was no strong evidence for publication bias in terms of an association between sample size and heritability estimate or genetic correlations, but there was evidence that both heritability estimates and genetic correlations decreased across time. These effects were independent of the other moderators including statistical methods that may also have changed across time, and potentially suggests that there is some time-lag bias, such that larger or more statistically significant results were published more quickly than smaller or non-significant results (Koricheva and Kulinskaya, 2019), and/or that newer studies have reduced error estimates due to larger sample sizes or more accurate estimation methods not captured in our meta-regression. Despite this effect, there was no evidence that accounting for this bias resulted in a large shift in the global estimate of either the heritability of resistance or its genetic correlation with performance.

This meta-analysis has provided a comprehensive, quantitative assessment of genetic parameters for resistance to GIN in sheep, and identified factors associated with variation in these parameters. Our first conclusion, that resistance to GIN has a heritable basis, is not surprising: decades of research show this to be the case, and have suggested a range of heritabilities for the trait that the global heritability estimated here lies within (Bishop and Morris, 2007; Saddiqi et al., 2011; Stear et al., 2009; Venturina et al., 2013; Windon, 1996). Importantly, however, this meta-analysis has revealed that this genetic basis of resistance is relatively stable: it does not differ substantially between breeds of sheep or with latitude, age, experimental design or statistical methods (**Table 1**). Furthermore, the main markers of parasite resistance that are used as a diagnostic and the basis of breeding (FEC, FAMACHA© and parasite-specific antibodies) all have moderate heritabilities. In addition, research shows that measures of parasite resistance are, in general, favourably correlated at the genetic level (Álvarez et al., 2018; Bishop et al., 2004; Davies et al., 2005; Douch et al., 1995a; Shaw et al., 2013). A bigger barrier to breeding for resistance has been the idea that a genetic trade-off may exist between resistance and other aspects of performance (Greer, 2008; Hoste and Torres-Acosta, 2011; Kloosterman et al., 1992), but this meta-analysis found no evidence that this would be the case when considering FEC and FAMACHA© scores (**Figure 2F**). On the other hand, it appears that breeding for parasite-specific antibody responses, specifically in the circulation, could lead to a negative response in performance traits. As with heritability estimates, genetic correlations did not vary consistently in studies of different breeds, latitudes, ages, or statistical methods, and there was also no variation among parasite species (**Table 2**). Overall, the results of this meta-analysis paint an encouraging picture and suggest that breeding for enhanced resistance should be a key tool in integrated parasite management programs alongside improved pasture management, nutrition, vaccines, and promotion of tolerance to infection (Hoste and Torres-Acosta, 2011; Jackson et al., 2009; Knap and Doeschl-Wilson, 2020; McManus et al., 2014; Sayers and Sweeney, 2005) in order to reduce anthelminthic use and improve the productivity and sustainability of the industry.

## ACKNOWLEDGEMENTS

I am greatly indebted to Thomas Connelly for his tireless efforts in tracking down obscure literature items; his work greatly enhanced the number of articles I was able to include. Sincere thanks also to Keith Ballingall, Tom McNeilly, Jocelyn Poissant and Alexandra Sparks for reading a draft of the manuscript and providing perceptive and constructive comment that improved it greatly. A number of other people also responded to my requests for articles and conference proceedings and took time to dig them out from places varying from hard drives to the garden shed: thank you to Leyden Baker, Doug Gray, Johan Greeff, Kathryn Kemper, Li Li, Thomas Pollott & Steve Walkden-Brown. I received funding from a Moredun Foundation Research Fellowship.

## DATA STATEMENT

Upon publication, all the data used in the study will be published online via the Dryad data repository.

## Notes

### Competing Interest Statement

The authors have declared no competing interest.

