## Supporting Information for "Genetic parameters for resistance to gastrointestinal nematodes in sheep: a meta-analysis"

Adam D. Hayward

**Table S1.** PRISMA checklist for the meta-analysis.

| **Topic** | **#** | **Checklist item** | **Reported where** |
| --- | --- | --- | --- |
| **TITLE** |  |  |  |
| **Title** | 1 | Identify the report as a systematic review, meta-analysis, or both. | P1 |
| **ABSTRACT** |  |  |  |
| **Structured summary** | 2 | Provide a structured summary including, as applicable: background; objectives; data sources; study eligibility criteria, participants, and interventions; study appraisal and synthesis methods; results; limitations; conclusions and implications of key findings; systematic review registration number. | P2 |
| **INTRODUCTION** |  |  |  |
| **Rationale** | 3 | Describe the rationale for the review in the context of what is already known. | P3-7 |
| **Objectives** | 4 | Provide an explicit statement of questions being addressed with reference to participants, interventions, comparisons, outcomes, and study design (PICOS). | P7 |
| **METHODS** |  |  |  |
| **Protocol and registration** | 5 | Indicate if a review protocol exists, if and where it can be accessed (e.g., Web address), and, if available, provide registration information including registration number. | NA |
| **Eligibility criteria** | 6 | Specify study characteristics (e.g., PICOS, length of follow-up) and report characteristics (e.g., years considered, language, publication status) used as criteria for eligibility, giving rationale. | P7-8 |
| **Information sources** | 7 | Describe all information sources (e.g., databases with dates of coverage, contact with study authors to identify additional studies) in the search and date last searched. | P7 |
| **Search** | 8 | Present full electronic search strategy for at least one database, including any limits used, such that it could be repeated. | P7 |
| **Study selection** | 9 | State the process for selecting studies (i.e., screening, eligibility, included in systematic review, and, if applicable, included in the meta-analysis). | P7-8; Tables S2&S3; Figure S1 |
| **Data collection process** | 10 | Describe method of data extraction from reports (e.g., piloted forms, independently, in duplicate) and any processes for obtaining and confirming data from investigators. | P8 |
| **Data items** | 11 | List and define all variables for which data were sought (e.g., PICOS, funding sources) and any assumptions and simplifications made. | P9-11 |
| **Risk of bias in individual studies** | 12 | Describe methods used for assessing risk of bias of individual studies (including specification of whether this was done at the study or outcome level), and how this information is to be used in any data synthesis. | P12-13 |
| **Summary measures** | 13 | State the principal summary measures (e.g., risk ratio, difference in means). | P9 |
| **Synthesis of results** | 14 | Describe the methods of handling data and combining results of studies, if done, including measures of consistency (e.g., I^2^) for each meta-analysis. | P11-12 |
| **Risk of bias across studies** | 15 | Specify any assessment of risk of bias that may affect the cumulative evidence (e.g., publication bias, selective reporting within studies). | P12-13 |
| **Topic** | **#** | **Checklist item** | **Reported where** |
| **Additional analyses** | 16 | Describe methods of additional analyses (e.g., sensitivity or subgroup analyses, meta-regression), if done, indicating which were pre-specified. | P12-13 |
| **RESULTS** |  |  |  |
| **Study selection** | 17 | Give numbers of studies screened, assessed for eligibility, and included in the review, with reasons for exclusions at each stage, ideally with a flow diagram. | P7-8; Fig S1 |
| **Study characteristics** | 18 | For each study, present characteristics for which data were extracted (e.g., study size, PICOS, follow-up period) and provide the citations. | Table S4 |
| **Risk of bias within studies** | 19 | Present data on risk of bias of each study and, if available, any outcome level assessment (see item 12). | P12-13 |
| **Results of individual studies** | 20 | For all outcomes considered (benefits or harms), present, for each study: (a) simple summary data for each intervention group (b) effect estimates and confidence intervals, ideally with a forest plot. | NA |
| **Synthesis of results** | 21 | Present results of each meta-analysis done, including confidence intervals and measures of consistency. | Tables 1 & 2; Figs 1 & 2 |
| **Risk of bias across studies** | 22 | Present results of any assessment of risk of bias across studies (see Item 15). | P14 & 17 |
| **Additional analysis** | 23 | Give results of additional analyses, if done (e.g., sensitivity or subgroup analyses, meta-regression [see Item 16]). | P13-19; Tables 1&2; Figs 1&2 |
| **DISCUSSION** |  |  |  |
| **Summary of evidence** | 24 | Summarize the main findings including the strength of evidence for each main outcome; consider their relevance to key groups (e.g., healthcare providers, users, and policy makers). | P20-26 |
| **Limitations** | 25 | Discuss limitations at study and outcome level (e.g., risk of bias), and at review-level (e.g., incomplete retrieval of identified research, reporting bias). | P20-26 |
| **Conclusions** | 26 | Provide a general interpretation of the results in the context of other evidence, and implications for future research. | P25-26 |
| **FUNDING** |  |  |  |
| **Funding** | 27 | Describe sources of funding for the systematic review and other support (e.g., supply of data); role of funders for the systematic review. | P26 |

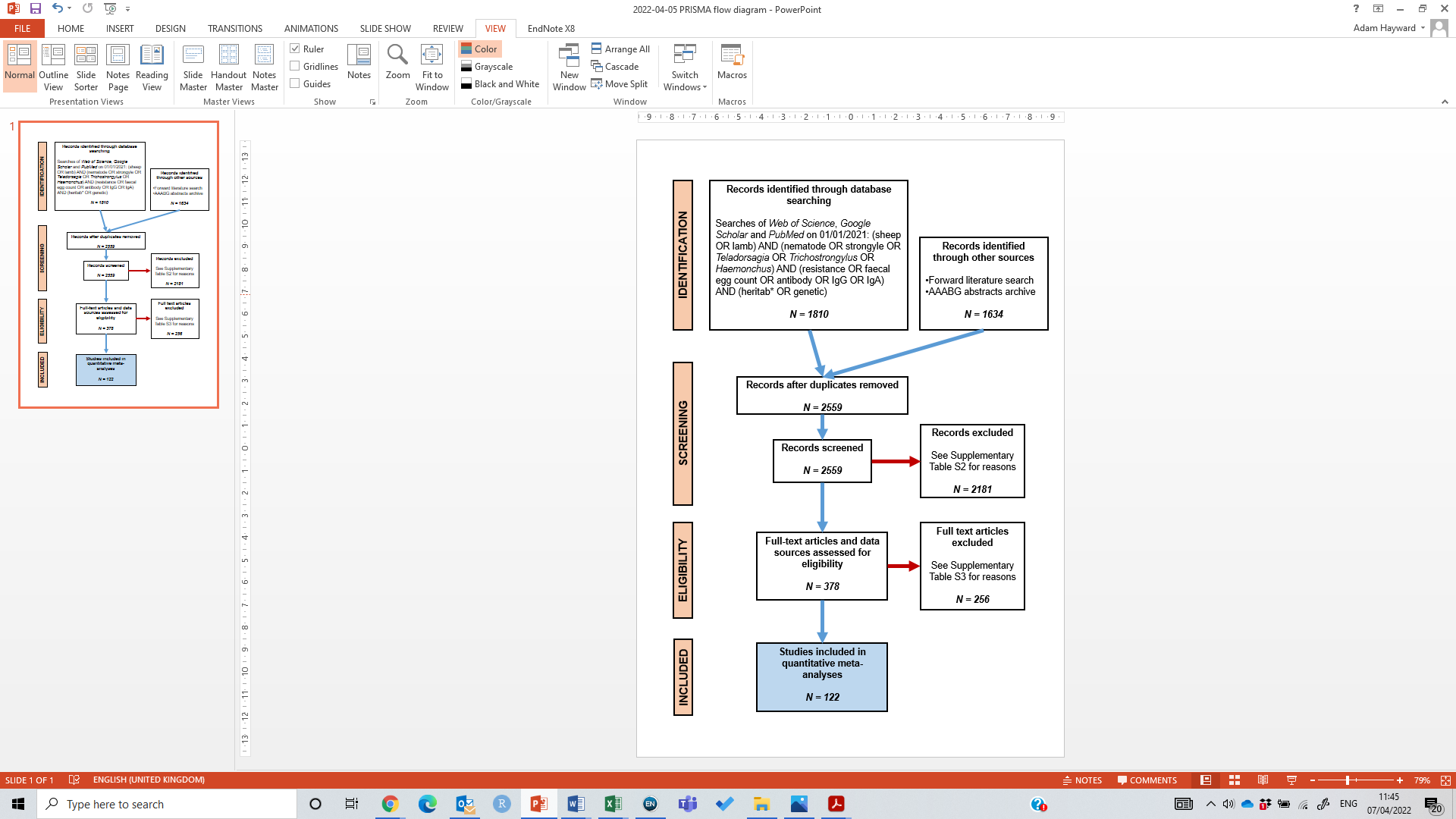

**Figure S1.** PRISMA flow diagram for the systematic meta-analysis.

**Table S2.** Results of the initial screen of 2,559 research items and reasons for their exclusion.

| **Reason** | **N** | **%** |
| --- | --- | --- |
| The study met inclusion criteria and was taken forward for full review | 378 | 14.77% |
| A study of diagnostic techniques | 28 | 1.09% |
| A study of MHC diversity and associations with performance | 29 | 1.13% |
| A gene expression study | 39 | 1.52% |
| A study not on gastrointestinal nematodes | 61 | 2.38% |
| A mathematical model not including empirical data | 66 | 2.58% |
| A study focused on genetic diversity of the parasite host | 141 | 5.51% |
| A study of the mechanisms of the immune response | 177 | 6.92% |
| A study of resistance phenotypes not including any genetic analysis | 179 | 6.99% |
| A study not including the traits of interest | 183 | 7.15% |
| A study comparing genotypes or breeds of sheep | 230 | 8.99% |
| A study of anthelmintic resistance in parasites | 342 | 13.36% |
| A study not on sheep | 351 | 13.72% |
| A study not suitable for a myriad of reasons | 355 | 13.87% |
| Total | 2559 | 100.00% |

**Table S3.** Results of the full screening of 378 research items and reasons for their exclusion from the final data set.

| **Reason** | **N** | **%** |
| --- | --- | --- |
| The study met inclusion criteria and provided data for analysis | 122 | 32.28% |
| A study of gene expression | 2 | 0.53% |
| A study not on sheep | 5 | 1.32% |
| A mathematical model with no empirical data | 6 | 1.59% |
| Studies that could not be tracked down | 7 | 1.85% |
| A study at the phenotypic level | 12 | 3.17% |
| A study reporting data repeated elsewhere | 13 | 3.44% |
| The data were not of suitable format for inclusion in the analysis | 14 | 3.70% |
| A genome-wide association study not providing heritability estimates | 15 | 3.97% |
| A study of the wrong trait or traits | 20 | 5.29% |
| The study was a duplicate | 24 | 6.35% |
| A study of quantitative trait loci | 25 | 6.61% |
| A study comparing performance of selection lines | 28 | 7.41% |
| A review article with no new empirical data | 85 | 22.49% |
| Total | 378 | 100.00% |

**Table S4.** The studies that provided data included in the meta-analyses of heritability and/or genetic correlations. Traits are FEC (faecal egg count), IMM (immune marker), FAM (FAMACHA score) and WO (worm traits); infection methods are N (natural) or C (challenge); analytical method is a pedigree-based ‘animal model’ or another method; parasites are ST (strongyle), HC (*Haemonchus contortus*) or NB (*Nematodirus battus*) dominant infections; traits are those included in genetic correlations and are NA (not applicable; study did not measure genetic correlations), BW (body weight), DWG (daily weight gain), W (wool traits), C (condition measure) or R (reproductive traits). N *h²* and N *r_g_* are the number of heritability and genetic correlation estimates provided by each study.

| **Reference** | **Country** | **Traits** | **Infection** | **Method** | **Parasites** | **Traits** | **N *h²*** | **N *r_g_*** |
| --- | --- | --- | --- | --- | --- | --- | --- | --- |
| Abuargob 2005 | UK | FEC, IMM | N | Animal | ST, NB | NA | 7 | 0 |
| Aguerre et al. 2018 | France | FEC | N, C | Animal | HC | NA | 3 | 0 |
| Al Kalaldeh et al. 2019 | Australia | FEC | N | Animal | HC | NA | 1 | 0 |
| Albers et al. 1987 | Australia | FEC | C | Other | HC | DWG, W, C | 2 | 12 |
| Álvarez et al. 2018 | Burkina Faso | FEC, FAM | N | Animal | HC | NA | 4 | 0 |
| Assenza et al. 2014 | France | FEC | C | Animal | HC | DWG | 4 | 8 |
| Baker et al. 1990 | NZ | FEC | N | Other | ST | DWG | 4 | 2 |
| Baker et al. 1994b | Kenya | FEC | N | Animal | HC | NA | 2 | 0 |
| Baker et al. 1994a | Kenya | FEC | N | Animal | HC | NA | 1 | 0 |
| Baker et al. 1998 | Kenya, Ethiopia | FEC | N | Animal | HC | NA | 9 | 0 |
| Baker et al. 2003 | Kenya | FEC | N | Animal | HC | BW | 24 | 6 |
| Benavides et al. 2016 | Brazil | FEC | N | Animal | HC | W | 1 | 8 |
| Beraldi et al. 2007 | UK | FEC | N | Animal | ST | NA | 2 | 0 |
| Berton et al. 2017 | Brazil | FEC, FAM | N | Animal | HC | NA | 2 | 0 |
| Berton et al. 2019 | Brazil | FEC, FAM | N | Animal | HC | BW, R | 0 | 6 |
| Bishop & Stear 2001 | UK | FEC | N | Animal | ST | R | 1 | 3 |
| Bishop et al. 1996 | UK | FEC | N | Animal | ST | BW | 6 | 2 |
| Bishop et al. 2004 | UK | FEC | N | Animal | ST, NB | BW, C | 16 | 36 |
| Bisset et al. 1992 | NZ | FEC | N | Other | ST, NB | BW, DWG, W | 2 | 4 |
| Bisset et al. 1994 | NZ | FEC | N | Other | ST | BW, DWG, W | 1 | 4 |
| Bisset et al. 2001 | South Africa | FEC, FAM | N | Animal | HC | BW, W | 2 | 6 |
| Bouix et al. 1998 | Poland | FEC | N | Animal | ST | DWG, R | 5 | 5 |
| Brown & Fogarty 2017 | Australia | FEC | N | Animal | ST | BW, W, C, R | 4 | 72 |
| Brown et al. 2013 | UK | FEC, IMM | N | Animal | ST | NA | 12 | 0 |
| Ceyhan et al. 2015 | UK | FEC | N | Animal | ST, NB | NA | 2 | 0 |
| Charon et al. 2000 | Poland | FEC | N | Animal | HC | DWG, W | 1 | 4 |
| Ciappesoni & Goldberg 2018 | Uruguay | FEC, FAM | N | Animal | HC | BW | 2 | 2 |
| Clement et al. 1999 | Senegal | FEC | N | Animal | ST | NA | 12 | 0 |
| Cloete et al. 2007 | South Africa | FEC | N | Animal | HC | BW, W | 3 | 5 |
| Cloete et al. 2016 | South Africa | FEC, FAM | N | Animal | HC | C | 4 | 8 |
| Coltman et al. 2001 | UK | FEC | N | Animal | ST | BW, S | 4 | 8 |
| Davies et al. 2005 | UK | FEC, IMM, WO | N | Animal | ST | NA | 17 | 0 |
| Davies et al. 2006 | UK | FEC, IMM | N | Animal | ST, NB | NA | 7 | 0 |
| Douch et al. 1995b | NZ | FEC, IMM | N | Animal | HC | BW, DWG, W | 7 | 28 |
| Douch et al. 1995a | NZ | FEC, IMM | N | Animal | ST | BW, DWG | 5 | 15 |
| Eady et al. 1996 | Australia | FEC | C | Other | ST, HC | NA | 7 | 0 |
| Eady et al. 1998 | Australia | FEC | C | Other | ST, HC | BW, W | 6 | 50 |
| Fairlie-Clarke et al. 2019 | UK | FEC, IMM | N | Animal | ST, NB | BW, C | 3 | 4 |
| Gauly & Erhardt 2001 | Germany | FEC | N | Other | ST | NA | 3 | 0 |
| Gauly et al. 2002 | Germany | FEC, IMM, WO | C | Other | HC | NA | 14 | 0 |
| Goldberg et al. 2012 | Uruguay | FEC | N | Animal | HC | NA | 2 | 0 |
| Gowane et al. 2019 | India | FEC | N | Animal | HC | BW | 4 | 8 |
| Gowane et al. 2020 | India | FEC | N | Animal | HC | NA | 4 | 0 |
| Graham et al. 2010 | UK | IMM | N | Animal | ST | NA | 10 | 0 |
| Greeff & Karlsson 1997 | Australia | FEC | N | Animal | ST | NA | 2 | 0 |
| Greeff et al. 2007 | Australia | FEC | N | Animal | ST | NA | 2 | 0 |
| Green et al. 1999 | NZ | FEC | N | Animal | ST | NA | 15 | 0 |
| Gruner et al. 2004b | France | FEC | C | Animal | ST | NA | 2 | 0 |
| Gruner et al. 2004a | France | FEC | C | Other | ST, HC | NA | 8 | 0 |
| Gutiérrez-Gil et al. 2010 | Spain | FEC, IMM | N | Animal | ST | NA | 3 | 0 |
| Haehling 2020 | Brazil | FEC | C | Animal | HC | BW, DWG | 16 | 8 |
| Hayward et al. 2014 | UK | IMM | N | Animal | ST | NA | 1 | 0 |
| Hollema et al. 2018 | Australia | FEC | N | Animal | ST | DWG | 3 | 3 |
| Keane et al. 2018 | Ireland | FEC | N | Animal | ST, NB | NA | 4 | 0 |
| Kemper et al. 2011 | Australia | FEC | C | Animal | ST, HC | NA | 2 | 0 |
| Khusro et al. 2004 | Australia | FEC | N | Animal | ST | BW, W | 2 | 8 |
| Li et al. 2015 | Australia | FEC | N | Other | ST | NA | 8 | 0 |
| Li et al. 2019 | Australia | FEC | N | Animal | ST | NA | 4 | 0 |
| Balconi Marques et al. 2020 | Uruguay | FEC, FAM | N | Animal | HC | BW, C | 4 | 8 |
| Matebesi-Ranthimo et al. 2014a | South Africa | FEC | N | Animal | ST | W | 1 | 6 |
| Matebesi-Ranthimo et al. 2014b | South Africa | FEC | N | Animal | ST | W, C | 0 | 15 |
| McEwan & Kerr 1998 | NZ | FEC | N | Animal | ST, NB | NA | 2 | 0 |
| McEwan et al. 1992 | NZ | FEC | N | Other | ST, NB | BW, DWG, W | 2 | 6 |
| McEwan et al. 1995 | NZ | FEC, IMM | N | Animal | ST, NB | NA | 10 | 0 |
| McManus et al. 2009 | Brazil | FEC | N | Animal | ST | NA | 2 | 0 |
| McRae 2015 | Ireland | FEC | N | Animal | ST, NB | NA | 4 | 0 |
| Miller et al. 2006 | USA | FEC | C | Other | ST | NA | 3 | 0 |
| Morris et al. 1998 | NZ | FEC | N | Animal | ST | NA | 1 | 0 |
| Morris et al. 2000 | NZ | FEC | N | Animal | ST | NA | 3 | 0 |
| Morris et al. 2004 | NZ | FEC | N | Animal | ST, NB | BW, DWG, W | 4 | 12 |
| Morris et al. 2005 | NZ | FEC | N | Animal | ST | BW, W | 3 | 2 |
| Morris et al. 2010 | NZ | FEC | N | Animal | ST | C | 1 | 1 |
| Mpetile et al. 2015 | South Africa | FEC | N | Animal | ST | NA | 1 | 0 |
| Mpetile et al. 2017 | South Africa | FEC | N | Animal | ST | NA | 3 | 0 |
| Murphy et al. 2010 | UK | IMM | N | Animal | ST | NA | 2 | 0 |
| Newton et al. 2011 | Australia | FEC | C | Animal | ST, HC | NA | 4 | 0 |
| Ngere et al. 2017 | South Africa | FEC, FAM | N | Animal | HC | NA | 8 | 0 |
| Ngere et al. 2018 | USA | FEC | N | Animal, Other | HC | BW, C | 10 | 6 |
| Nieuwoudt et al. 2002 | South Africa | FEC | N | Animal | HC | BW | 1 | 1 |
| Notter et al. 2018 | USA | FEC | N | Animal | HC | R | 2 | 10 |
| Oliveira et al. 2018 | Brazil | FEC, FAM | N | Animal | HC | BW, C | 2 | 4 |
| Pacheco et al. 2021 | UK | FEC | N | Animal | ST, NB | BW | 2 | 2 |
| Pfeffer et al. 2007 | NZ | FEC | N | Animal | ST | NA | 3 | 0 |
| Pickering et al. 2015 | NZ | FEC | N | Animal | ST | NA | 2 | 0 |
| Piper 1987 | Australia | FEC | C | Other | HC | BW, W, R | 1 | 5 |
| Pollott & Greeff 2004 | Australia | FEC | N | Animal | ST | BW, W, C | 2 | 16 |
| Pollott et al. 2004 | Australia | FEC | N | Other | ST | BW | 9 | 9 |
| Prince et al. 2010 | India | FEC | N | Animal | HC | NA | 1 | 0 |
| Rashidi 2016 | UK | FEC, IMM | N | Other | ST | NA | 2 | 0 |
| Rice et al. 1992 | Australia | FEC, IMM | C | Other | ST, HC | NA | 5 | 0 |
| Riggio et al. 2013 | UK | FEC, IMM | N | Animal | ST, NB | NA | 7 | 0 |
| Riley & Van Wyk 2009 | South Africa | FEC, FAM | N | Animal | HC | BW, DWG, C | 3 | 6 |
| Rodrigues et al. 2021 | Brazil | FEC, FAM | N | Animal | HC | BW, C | 2 | 4 |
| Roy et al. 2018 | India | FEC | N | Other | HC | NA | 12 | 0 |
| Salle 2010 | UK | FEC | N | Animal | ST | BW, C | 6 | 18 |
| Sechi et al. 2009 | Italy | FEC | N | Animal | ST | NA | 1 | 0 |
| Shaw et al. 1999 | NZ | FEC, IMM | N | Animal | ST | BW, DWG | 9 | 27 |
| Shaw et al. 2012 | NZ | FEC, IMM | N | Animal | ST | NA | 6 | 0 |
| Shaw et al. 2013 | NZ | FEC, IMM | N | Animal | ST | BW | 7 | 36 |
| Singh et al. 1999 | India | FEC | N | Other | HC | NA | 1 | 0 |
| Smith et al. 1999 | UK | FEC | N | Other | ST | NA | 21 | 0 |
| Snyman & Fisher 2019 | South Africa | FEC, FAM | N | Animal | HC | C | 24 | 2 |
| Sparks 2018 | UK | IMM | N | Animal | ST | NA | 7 | 0 |
| Sparks et al. 2019 | UK | IMM | N | Animal | ST | NA | 6 | 0 |
| Sréter et al. 1994 | Australia | FEC | C | Other | HC | NA | 1 | 0 |
| Stear et al. 1997 | UK | FEC, WO | N | Animal | ST | NA | 6 | 0 |
| Stear et al. 2002 | UK | IMM | N | Animal | ST | NA | 2 | 0 |
| Strain et al. 2002 | UK | IMM | N | Animal | ST | NA | 1 | 0 |
| Swarnkar et al. 2009 | India | FEC | N | Other | HC | NA | 2 | 0 |
| Torres et al. 2021 | Brazil | IMM | N | Animal | HC | S | 2 | 4 |
| Vagenas et al. 2007 | UK | FEC | N | Animal | ST | NA | 5 | 0 |
| Vanimisetti 2003 | USA | FEC | C | Animal | HC | NA | 2 | 0 |
| Watson et al. 1986 | NZ | FEC | N | Other | ST, NB | NA | 7 | 0 |
| Windon 1985 | Australia | FEC | C | Other, Other | ST | NA | 9 | 0 |
| Windon et al. 1988 | Australia | FEC | C | Other | ST | NA | 1 | 0 |
| Wolf et al. 2008 | UK | FEC | N | Animal | ST, NB | BW, C | 8 | 20 |
| Woolaston & Piper 1996 | Australia | FEC | C | Animal | HC | NA | 1 | 0 |
| Woolaston & Windon 2001 | Australia | FEC | C | Animal | ST | NA | 11 | 0 |
| Woolaston et al. 1990 | Australia | FEC | C | Animal | ST, HC | NA | 4 | 0 |
| Woolaston et al. 1996 | Australia, Fiji | FEC, IMM | N, C | Animal | ST | NA | 4 | 0 |
| Yadav et al. 2006 | India | FEC | N | Other | HC | BW | 1 | 1 |
| Zhao et al. 2019 | Australia | IMM | N | Animal | ST | NA | 6 | 0 |

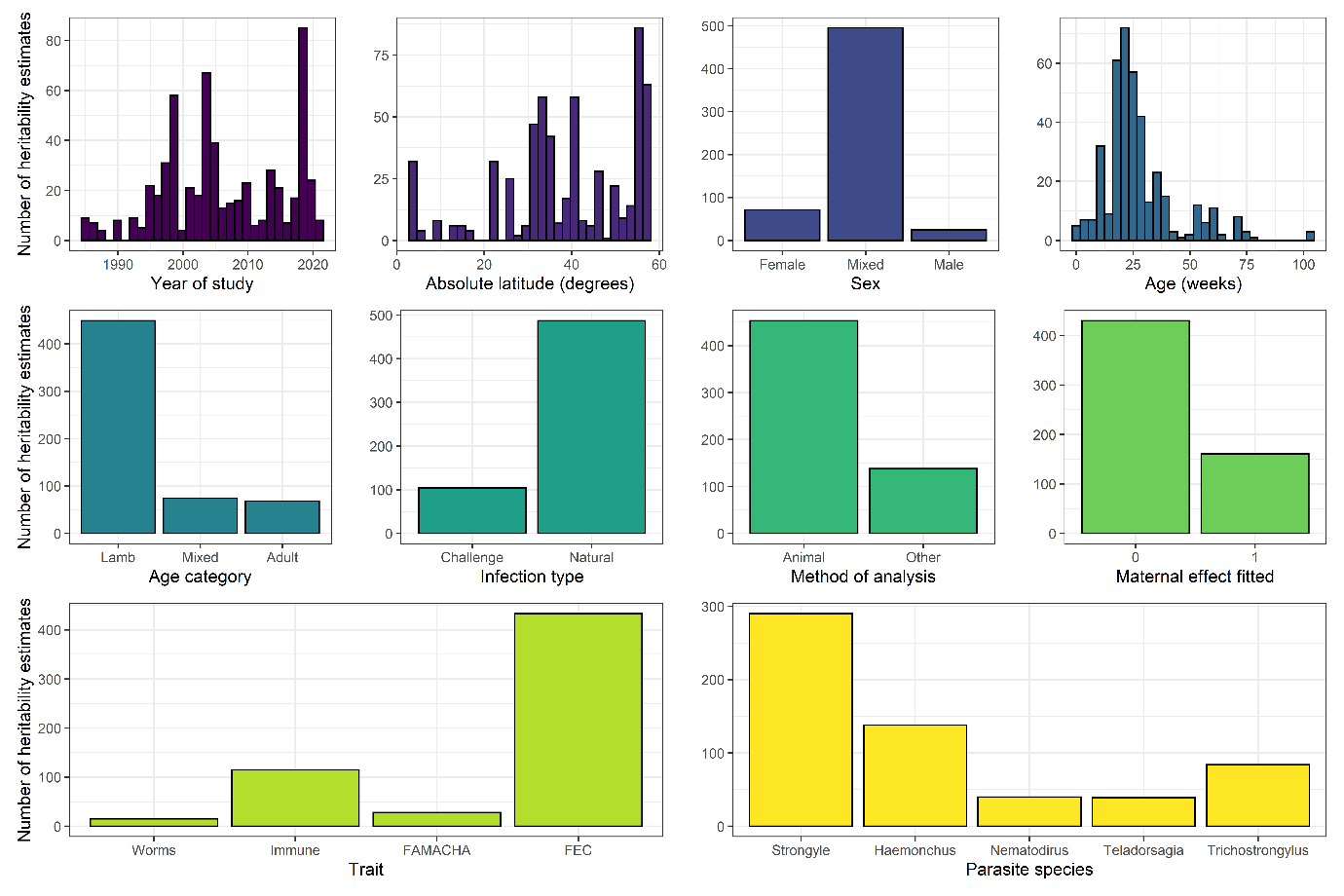

**Figure S2.** A breakdown of heritability estimates by moderator variables used in meta-regression analysis. In the ‘Method of analysis’ plot, ‘Animal’ indicates analysis by ‘animal model’.

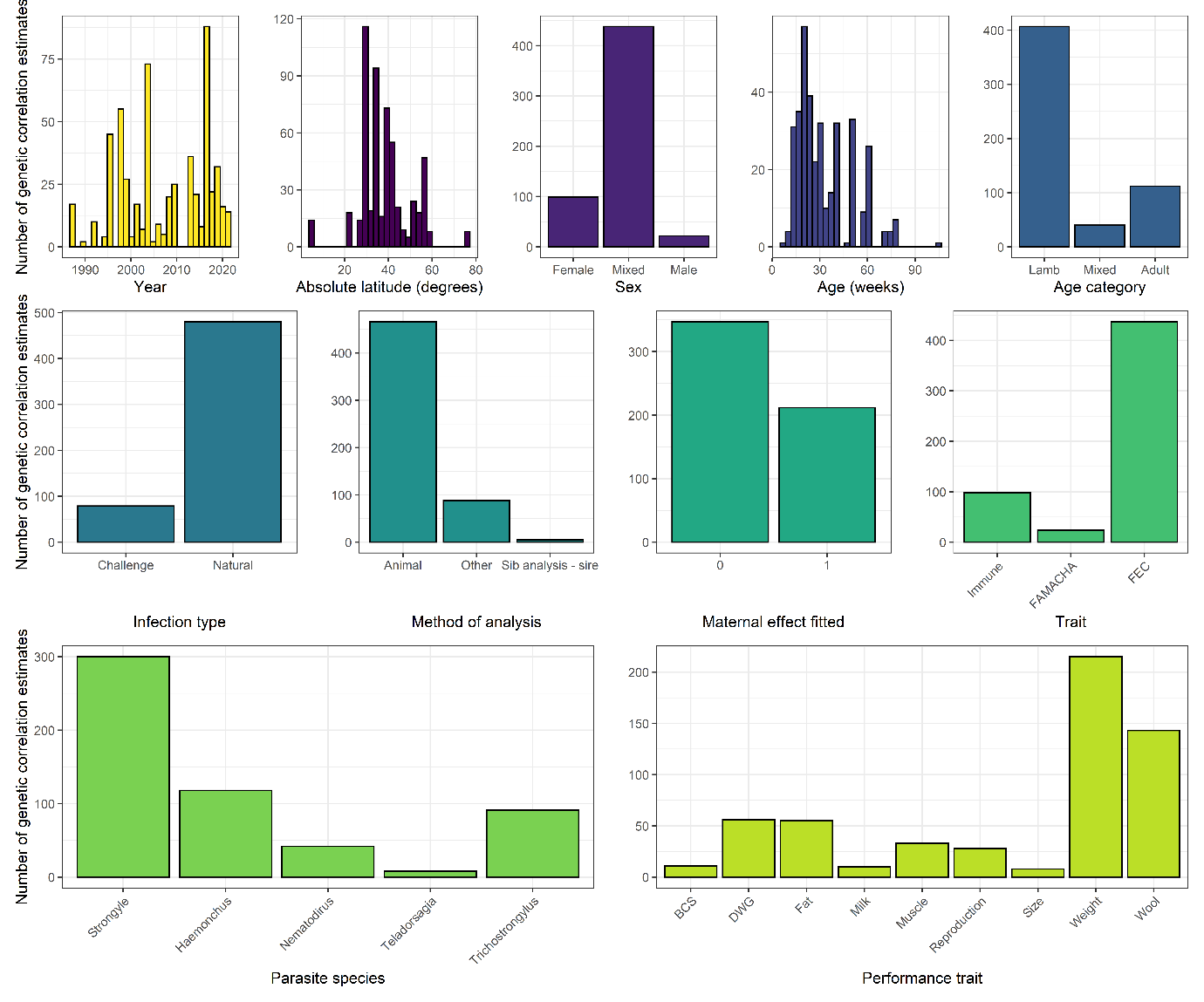

**Figure S3.** A breakdown of genetic correlations estimates by moderator variables used in meta-regression analysis.
